# It’s about time: neural temporal scaling accounts for robust hunting behavior across temperatures

**DOI:** 10.1101/2025.07.20.665717

**Authors:** Shai Tishby Tamari, Yoav Rubinstein, Netta Livneh, Maayan Moshkovitz, Abeer Karmi, Lilach Avitan

## Abstract

Animals are often required to maintain stable performance in critical behaviors despite environmental fluctuations. Temperature broadly affects neural activity, and even localized shifts in brain temperature can alter behavior. However, whether widespread changes across the brain, such as those experienced by ectotherms, disrupt survival-critical behaviors remains unclear. Here, we show that larval zebrafish maintain robust hunting performance across a 10°*C* ecological range. Although behavior accelerates with temperature, spatial parameters, such as bout distance and turn angle, remain stable. This invariance results from coordinated adjustments in tail beat frequency and movement duration. Brain-wide calcium imaging revealed that behavioral temporal scaling is mirrored at the level of single neurons. A simple rate model showed that temperature-dependent changes in neural time constants can account for compensatory tail dynamics, enabling stability without active regulation. These findings suggest that neural temporal scaling can preserve performance under diffuse temperature fluctuations, supporting robust behavior in natural environments.

## Introduction

To survive, animals often need to perform critical behaviors, such as escaping predators, capturing prey, or engaging in social interactions, even as environmental conditions change. Maintaining performance across such conditions, animals rely on adaptive mechanisms that range from active adjustments based on sensory input to intrinsic circuit properties that support robustness without requiring explicit control. In particular, some adaptations involve sensing environmental changes and modulating motor output accordingly ^1,2,3,4^, while others arise from self-organizing dynamics within neural circuits ^5,6,7^. Identifying these circuit-level forms of adaptation, active or self-organizing, remains a key challenge, as neural changes typically occur at the microscale, while their behavioral consequences emerge at the macroscale. While various approaches have been used to link these scales, one powerful strategy is brain-wide, single-cell resolution imaging during behavior, which enables direct observation of distributed neural dynamics.

Temperature is an environmental variable that modulates both behavior and neural activity across species^1^. Animals employ different behavioral strategies for thermoregulation, such as forming groups to conserve heat ^2,8^ or seeking sun exposure to increase body temperature ^2,3,9^. Temperature also directly affects neural dynamics ^1,10,11^, with immediate effects on behavior. For instance, in rodents, local cooling of the basal ganglia alters time perception ^12^, and in songbirds, manipulating the temperature of the premotor nucleus HVC shifts vocal timing ^13,14^. While these findings highlight temperature sensitivity of neural circuits, they typically rely on localized temperature manipulations that do not reflect ecological conditions.

Ectothermic animals, by contrast, naturally experience global fluctuations in brain and body temperature. This raises a fundamental question: can ectotherms maintain high-precision, survival-critical behaviors under thermal perturbations? The evidence to date is mixed. Some neural circuits, such as crustacean central pattern generators, maintain stable rhythmic output across a wide temperature range^6,15^. Conversely, other systems show clear temperature sensitivity; for example, prey-detecting neurons in dragonflies exhibit temperature-dependent changes in response latency and gain ^16^, and similar effects have been observed in goldfish, where cold temperatures lead to delayed neural motor responses and altered response gain ^17^. However, whether such neural changes result in behaviorally meaningful instabilities that compromise survival remains unclear. More broadly, it is still unknown whether ectotherms can achieve behavioral stability under global brain temperature shifts, and if so, what circuit-level mechanisms might support such robustness.

Zebrafish offer a powerful model for studying the neural basis of behavioral adaptation to temperature. They inhabit environments with wide thermal variability (18–34°*C*) ^18,19^ but must reliably perform survival-critical behaviors such as hunting, which demand precise timing and coordination ^20,21,22,23^. Larval zebrafish are optically transparent, enabling brain-wide, single-cell imaging during behavior, and providing a unique opportunity to link neural dynamics to natural behavior. Active thermoregulation guides their behavior in thermal gradients, with fish preferring 27–28°*C* and adjusting their position accordingly ^4,24,25,26^. At constant ecological temperatures, zebrafish modulate their spatial parameters of movement during exploratory behavior, with higher temperatures resulting in longer travel distances and sharper turns^27,28^. These movements occur as discrete episodes, termed bouts, consisting of brief, rapid swims separated by pauses. While temperature-dependent modulation of bouts is well-characterized during exploration, it remains unknown whether similar modulation extends to survival-critical behaviors like hunting. If it does, one would expect compensatory changes in hunting strategy to preserve performance. Alternatively, spatial movement parameters may remain stable, suggesting a neural mechanism underlying the robustness to thermal variation. In the absence of such compensation, performance may degrade outside the preferred thermal range.

Here, we examined how temperature affects zebrafish hunting behavior and the underlying neural dynamics. We found that while exploratory behavior changed with temperature, hunting performance remained stable across a 10°*C* range. This behavioral stability was accompanied by temperature-dependent temporal scaling: as temperature increased, bout durations, inter-bout intervals (IBIs), and total hunt duration became shorter. Despite these changes, fish maintained a stable distance traveled through coordinated modulation of tail beat frequency and movement duration. Brain-wide imaging across a similar temperature range revealed that temporal scaling extended to the activity of single neurons. A simple rate model demonstrated that temperature-dependent changes in time constants of individual neurons were sufficient to produce the observed coordination of tail beat frequency and duration to maintain similar bout distance during hunt, without requiring active compensation. Together, these findings reveal a fundamental circuit-level mechanism that supports behavioral stability in dynamic environments.

## Results

### Temperature modulates spatial movement statistics during exploration but not during hunting

To examine how temperature affects larval zebrafish behavior, we recorded 13 days post fertilization (dpf) larvae hunting Paramecia under three ecological temperature conditions: cold (22°*C*), intermediate (27°*C*), and hot (32°*C*) (*n* = 30, 10 fish per condition, Figure 1A, Supplementary Movie S1, see Methods). We selected this developmental stage because by 13 dpf, hunting behavior is fast, accurate, and robust, coinciding with the refinement of the tectal neural code compared to earlier stages, such as 7 dpf ^29^. Each fish was acclimated to the test temperature for at least 20 minutes before being placed in a plate containing Paramecia. A custom tracking system extracted frame-by-frame features including fish contour, heading direction, tail and eye postures, and prey positions (Figure 1B, see Methods). Hunting events were automatically detected based on eye convergence ^30,31^ (Figure S1A, see Methods), and individual movement bouts were segmented using tail posture dynamics (Figure S1B, see Methods). Hunting events were naturally interspersed with exploratory segments, during which the fish swam freely without targeting prey (Figure 1C, Figure S1C). This experimental design enabled us to assess temperature-dependent changes in behavior in the presence of prey. We compared spatial features of movement bouts, such as distance traveled and change in heading angle following each movement, and temporal features of movement bouts, such as bout and inter-bout interval (IBI) durations. This also allowed us to examine how hunting strategies adapt across temperature conditions.

**Figure 1:**
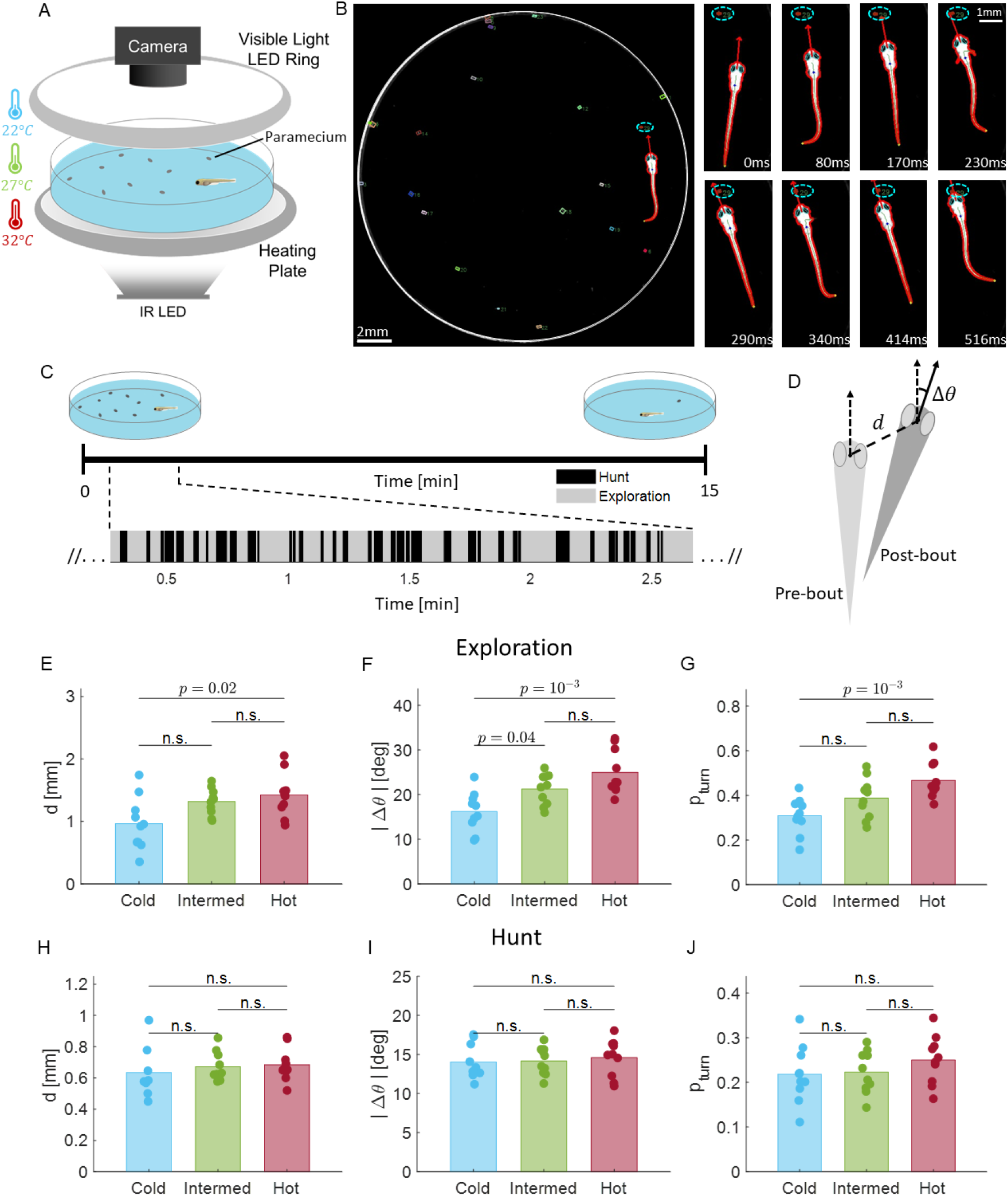
Temperature modulates larval zebrafish spatial movement statistics during exploration but not during hunt. **(A)** A schematic of the behavioral setup. A larval zebrafish was placed in a dish with 30 Paramecia, illuminated with visible and infrared light, and recorded for 15 minutes (500 fps). The temperature was controlled and monitored under three different conditions using a heating plate and a thermocouple sensor, respectively. **(B)** Left: an example annotated frame from a hunting event, showing the detected fish with its heading direction (red contour and red arrow, respectively), and Paramecia (colored objects). The target Paramecium is marked with a cyan ellipse. Right: a zoomed-in sequence of frames illustrating hunting event progression. The sequence includes the frame prior to eye convergence (i.e., event onset, 0 ms), initial turn and eye convergence (80 ms), approach movements and their respective IBIs (170–414 ms), and a strike movement (516 ms). **(C)** Hunting events (black) were interspersed with exploratory segments (gray). **(D)** Two measured spatial movement parameters: the distance traveled (*d*), and the change in heading angle (Δ*θ*) following a bout. Dashed and solid arrows indicate the pre- and post-bout heading direction, respectively. **(E)-(G)** During exploration, (E) bout distance was greater in the hot condition compared to the cold condition. (F) The absolute change in heading angle was larger in the hot compared to the cold condition.Turn probability (|Δ*θ*| *>* 20°) was higher in the hot condition compared to the cold condition. **(H)-(J)** During hunt, (H) bout distance did not differ across temperatures. (I) Absolute changes in heading angle did not differ across temperatures. (J) Turn probability did not differ across temperatures. Throughout the panels, each point indicates the mean value for a single fish. Temperature comparisons were performed using ANOVA, with Tukey-Kramer corrected p-values noted on all panels.

Temperature has been shown to modulate the spatial features of exploratory behavior in zebrafish, increasing travel distance and turn angle at higher temperatures and reducing them at lower ones ^27^. We asked whether this modulation persists during exploratory segments in the presence of prey, and whether a similar modulation extends to hunting behavior, potentially impacting hunting performance or strategy. For both exploratory and hunting movements, we calculated the distance traveled (*d*) and the change in heading angle (Δ*θ*) following each single movement bout (Figure 1D). During exploratory segments between hunts, fish in hot temperature traveled longer distances than those in cold temperature (Figure 1E, Figure S1D), and changes in heading angle increased with temperature (Figure 1F, Figure S1E). Turn probability was also higher in the hot condition compared to the cold condition (Figure 1G). These findings confirm that previously reported temperature-dependent modulation of exploratory behavior persists even in the presence of prey.

Given that exploratory spatial movement statistics were temperature-dependent, we next asked whether a similar modulation occurs during hunting, where movement precision is crucial. Interestingly, during hunting events, bout distance remained stable across temperatures (Figure 1H, S1F), as did the change in heading angle and turn probability (Figures 1I,J, Figure S1G). A significant interaction between temperature and behavioral context (exploration versus hunting) was observed for all movement metrics (two-way ANOVA interaction effects: *d*: *p* = 0.04;|Δ*θ*|: *p* = 0.003; p_*turn*_: *p* = 0.03), indicating that movement spatial statistics are differentially regulated by temperature depending on context. Together, while temperature modulates exploratory behavior, zebrafish maintain stable movement spatial statistics during hunting, potentially to preserve behavioral performance under changing environmental conditions. Therefore, we focused on how fish maintain this behavioral stability across temperatures during the hunt.

### Hunting behavior temporally scales with temperature

In prey-free environments, exploring fish show shorter IBI duration as temperature increases^27^. Similarly, during exploratory segments between hunts, IBI duration decreased with increasing temperature (Figure S2A). Interestingly, not only did the intervals shorten with temperature, but we found that bout duration during exploration was also shorter with the increase in temperature (Figure S2B). We then asked whether movements during hunting events show similar temperature-dependent temporal modulation as exploratory movements, or whether, like bout distance and change in heading angle, they remain temporally stable across temperatures. To address this, we pooled hunting events across all fish within each temperature condition, aligned them to the onset of the first bout, and sorted them by event duration (Figure 2A). Hunting events were shorter at higher temperature and longer at lower temperature, appearing “condensed” in the hot condition and “stretched” in the cold one (Figure 2B). This temporal scaling was evident in both bout duration (Figure 2C) and IBI duration (Figure 2D). Importantly, hunting behavior accelerated with temperature even when prey velocity was similar across temperature conditions (Figure S2C-I, see Methods), suggesting that the observed temporal scaling arises primarily from temperature effects on the fish rather than on prey motion. Together, these results indicate that temperature-dependent temporal scaling of hunting behavior, from individual movements to the full hunt sequence, occurs while preserving stable spatial movement statistics.

**Figure 2:**
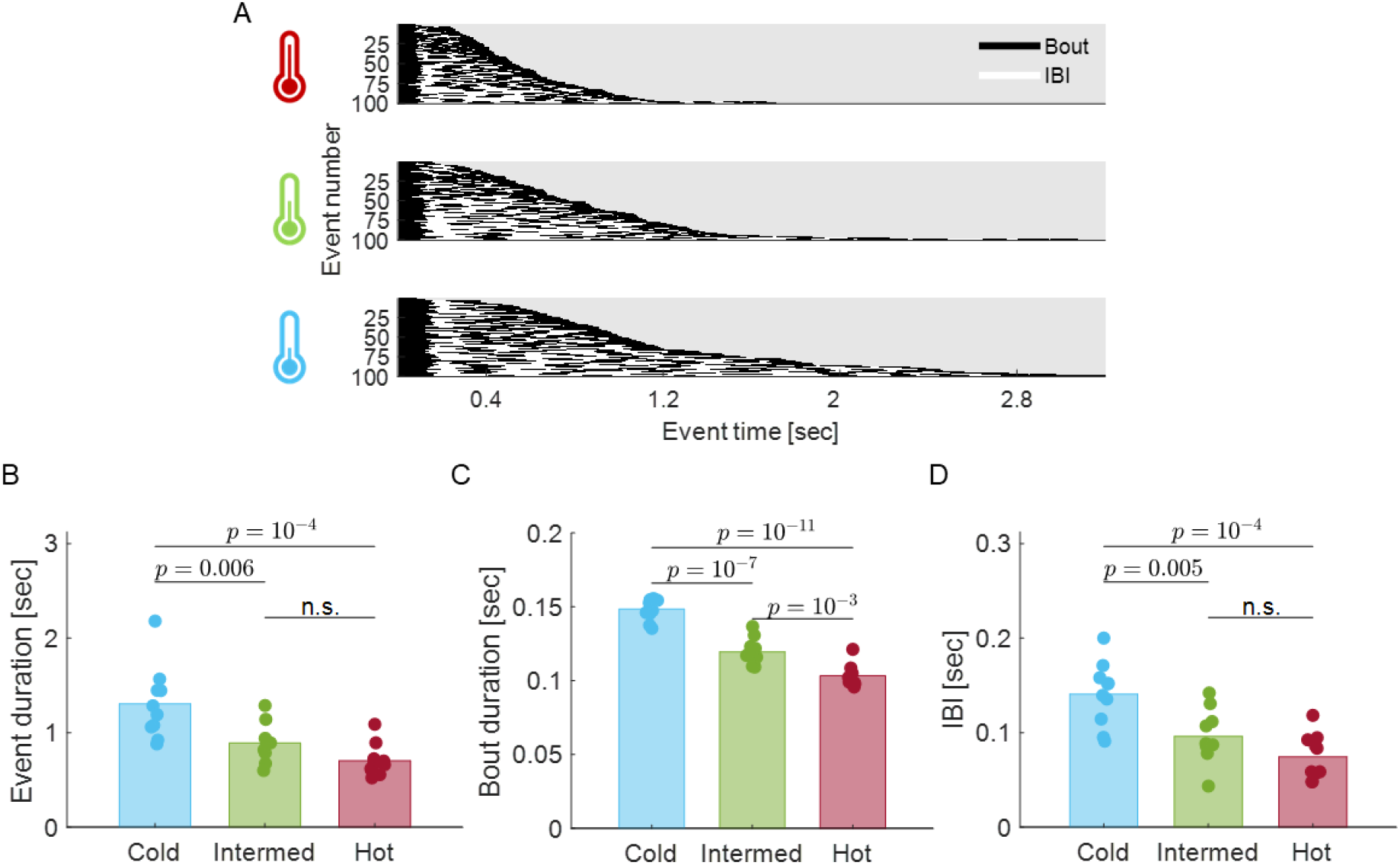
Temperature-dependent temporal scaling of hunting behavior. **(A)** 100 randomly selected successful hunting events from each temperature condition, sorted by event duration. Events were aligned by the onset of the first movement of the event. Black and white lines indicate time points of bouts and IBIs, respectively. A “condensing” effect was observed as temperature increased. **(B)** Hunting event duration was longer in the cold condition compared to the intermediate and hot conditions. **(C)** Bout duration decreased with increasing temperature. **(D)** IBI duration was longer in the cold condition compared to the intermediate and hot conditions.

### Hunting performance and strategy are preserved across temperatures

Since an increase in temperature shortened IBI and movement durations, we asked whether these changes impact hunting performance or strategy, potentially reflecting a trade-off between speed and accuracy. Hunting success rate, measured as the hit ratio, remained stable across temperatures (Figure 3A). To assess potential changes in strategy, we examined how fish approached their prey. Neither fish trajectory length during hunting events (Figure 3B), nor the number of bouts per event (Figure 3C) differed significantly across temperatures. Additionally, previous studies have reported a linear relationship between prey angle relative to the fish (*ϕ*_*prey*_) and the reactive change in heading angle (Δ*θ*) (Figure 3D, inset) ^23,32,33^. We found that this relationship remained consistent across temperatures (Figures 3D-F, permutation test *p* = 0.1). Together, these results indicate that fish preserve prey interaction strategies and robust hunting performance across temperatures, despite temperature-dependent temporal scaling.

**Figure 3:**
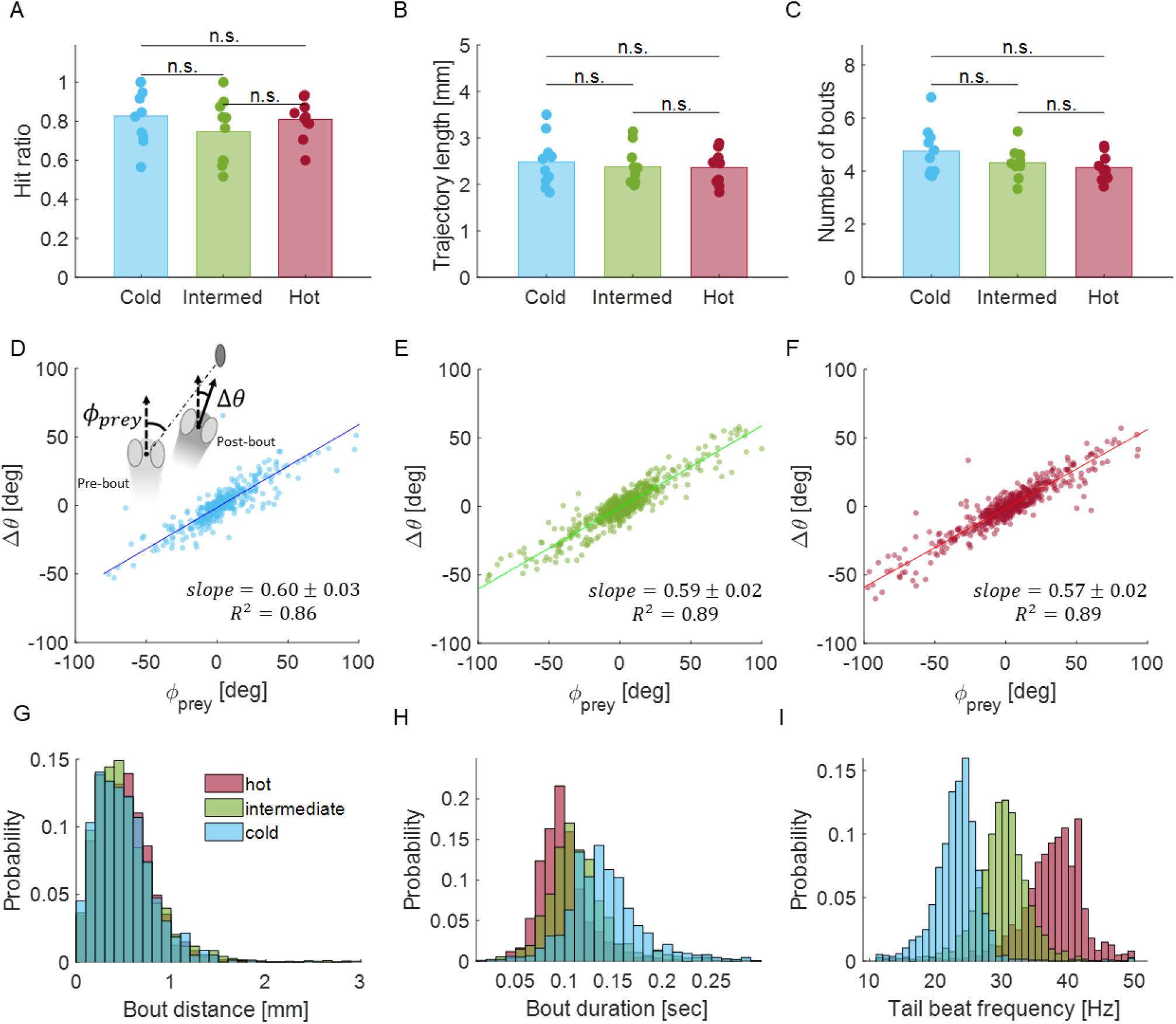
Coordinated modulation of movement duration and tail beat frequency allow behavioral stability across temperatures. **(A)** Hit ratio, measured as the fraction of events in which fish successfully struck and captured prey, remained consistent across temperatures. **(B)** Fish trajectory length during hunting events, from eye convergence to prey capture, did not differ across temperatures. **(C)** The number of bouts per hunting event did not differ across temperatures. **(D-F)** D: Inset: a schematic illustration of the prey angle before movement (*ϕ*_*prey*_) and the reactive change in heading angle (Δ*θ*). The pre- and post-bout fish heading direction are indicated by dashed and solid arrows, respectively. The relationship between prey angle (*ϕ*_*prey*_) and the reactive change in heading angle (Δ*θ*) remained consistent across temperature conditions (D: cold, E: intermediate, F: hot; least squares linear fit). **(G)** Bout distance distributions of forward bouts are similar across temperature conditions (Bonferroni-corrected KS-test, hot vs. intermediate *p* = 0.21, intermediate vs. cold *p* = 1, hot vs. cold *p* = 0.91).Bout duration distributions differed significantly across temperatures (Bonferroni-corrected KS-test, hot vs. intermediate *p* = 10^−44^, intermediate vs. cold *p* = 10^−89^, hot vs. cold *p* = 10^−193^). **(I)** Tail beat frequency differed significantly across temperatures (Bonferroni-corrected KS-test, hot vs. intermediate *p* = 10^−310^, intermediate vs. cold *p* = 0, hot vs. cold *p* = 0).

We next asked how fish maintain stable travel distances during hunting across temperatures (Figure 1H) despite temperature-dependent differences in movement duration (Figure 2C). To address this, we compared similar movements across temperatures, and focused on forward bouts (Δ*θ* < 20°), which accounted for 76.9% of all hunting bouts across conditions (Figure S3A). While the distance traveled during these bouts remained stable across temperatures (Figure 3G), their duration varied, being shortest in the hot condition and longest in the cold condition (Figure 3H). We hypothesized that this distance stability was achieved through adjustments of tail dynamics, specifically by coordinated modulation of movement duration and tail beat frequency, which correlates with swim velocity ^34,35,36^. Using Fourier analysis, we estimated the tail beat frequency of each movement (see Methods). Tail beat frequency increased with temperature (Figure 3I, Figure S3B), and was inversely correlated with bout duration (Figure S3C). These results suggest that both bout duration and tail beat frequency are jointly adjusted to maintain a consistent bout distance across temperatures during hunting behavior.

### Tectal neural coding of prey-like stimuli is robust to temperature

Since hunting behavior remained stable despite temperature-driven temporal scaling, we next asked whether neural processing of visual stimuli exhibited similar robustness. To test this, we simultaneously recorded neural activity and tail movements of 14 dpf larval zebrafish that were head-fixed and tail-free, presented with prey-like stimuli (Figure 4Ai). Each fish (n=14) was recorded in three temperatures: cold (23°*C*), intermediate (27.5°*C*), and hot (31.5°*C*) (Figure 4Aii, see Methods). We performed two-photon calcium imaging to record activity from a single brain plane (30 fps), focusing initially on the optic tectum, a key brain region for visual processing during hunt ^29,31,37,38,39^(Figure 4Aiii, Supplementary Movie S2). Simultaneously, we recorded tail movements (500 fps) to capture behavior with high temporal precision (Figure 4Aiv, Supplementary Movie S3). In each temperature condition, fish were shown a pseudo-random sequence of stationary (−60°,−30°, 30°, 60° relative to fish midline) and moving (−55°→ −10° and 55°→ 10°) prey-like stimuli (Figure 4v, see Methods). Overall, at each temperature, we collected responses to 80 stationary and 30 moving stimuli per fish.

**Figure 4:**
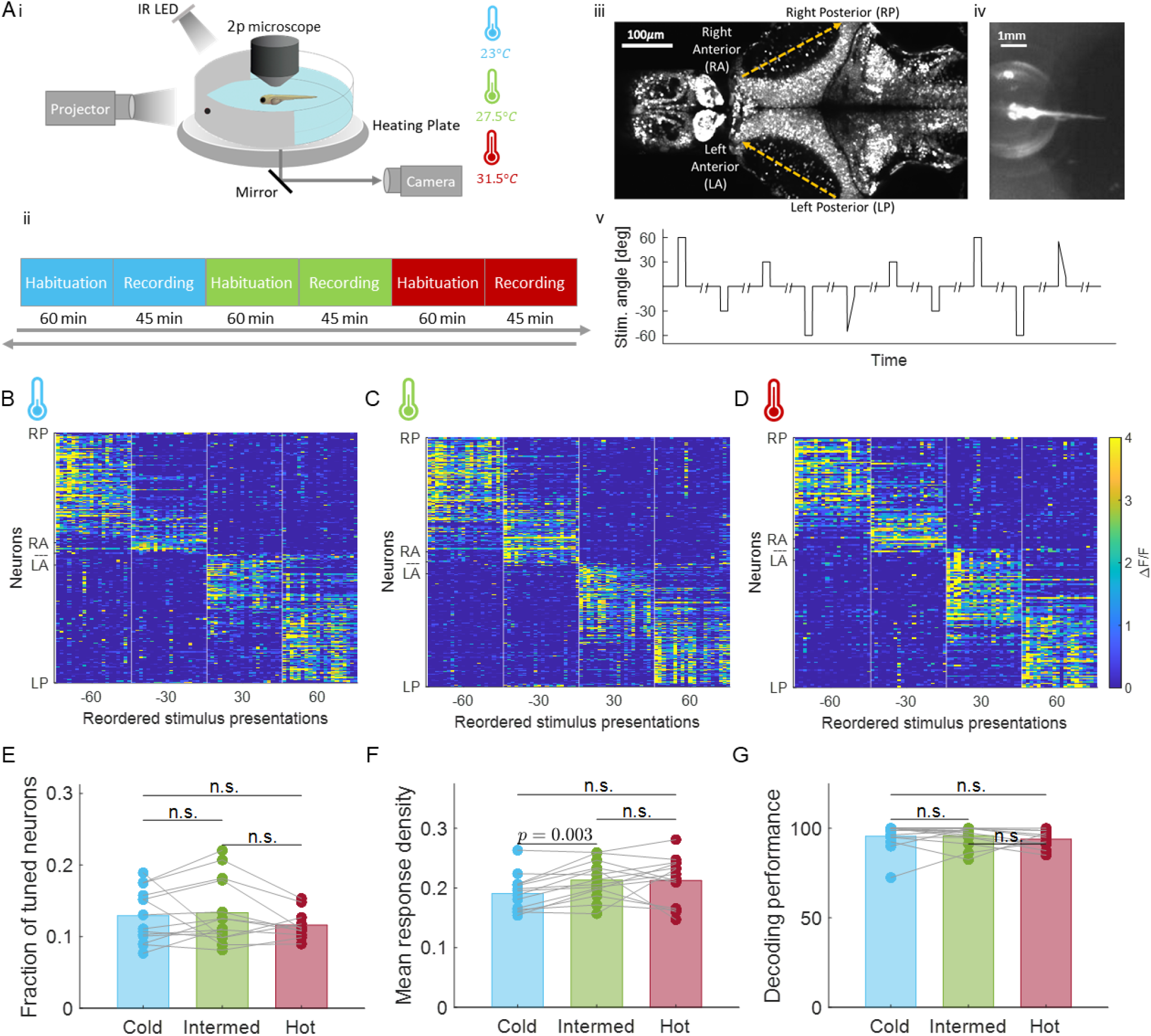
Tectal sensory neural responses remain stable across temperatures. **(A)** i. A schematic illustration of the experimental setup: a single 14 dpf larval zebrafish was embedded in a plate and placed on a heating plate under a two-photon microscope for a single-plane calcium imaging. Tail movements were simultaneously recorded from below with a high-speed camera. ii. Fish were habituated to each temperature condition for 1 hour, starting from either cold (n=7, rightward gray arrow) or hot (n=7, leftward gray arrow) temperature, followed by a 45-minute recording session at each temperature. iii. A mean image over 1000 recorded frames from two-photon calcium imaging. Tectal anterior-posterior (AP) axes (dashed yellow arrows) were used to order tectal neurons along these axes (see Methods). Right anterior and posterior (RA, RP) and left anterior and posterior (LA, LP) ends of the tectum are labeled. iv. An example frame from behavioral recording during two-photon calcium imaging. v. Visual stimuli presentation: stationary stimuli in four angular positions and moving stimuli in two directions were presented to the fish in a pseudo-random order. **(B)-(D)** Tectal population responses across temperatures showing a topographical organization, example from a single fish. Neurons were arranged according to their position on the AP axes, and responses were sorted by stimulus. (B) Cold, (C) Intermediate, (D) Hot conditions. **(E)** Fraction of tuned neurons remained stable across temperatures. **(F)** Tectal response density was lower for the cold condition compared to the intermediate condition but similar between the hot condition and the other two. **(G)** Decoding performance remained high and consistent across temperatures. Each point indicates the mean value for a single fish in a single temperature condition (indicated by color). Connected lines indicate recordings of the same fish. Temperature comparisons were performed using repeated-measures ANOVA, with Tukey-Kramer corrected p-values noted on the panels.

We identified tectal neurons tuned to prey-like stimuli using regressor-based analysis (Figure S4A, see Methods). These tuned neurons were consistently recruited within tectal population responses that displayed topographical organization across temperatures (Figures 4B-D, example from a single fish; Figure S4B for all fish). The proportion of tuned neurons per fish did not differ across temperatures (Figure 4E). Population response density, defined as the fraction of tuned neurons recruited per stimulus presentation, was slightly lower in the cold condition compared to the intermediate condition, but similar between the hot and both cold and intermediate conditions (Figure 4F). Population response amplitude followed a similar pattern, with a mild reduction in the cold condition, but similar levels in hot and intermediate conditions (Figure S4C). Given this overall stability, we asked whether tectal neural activity could reliably decode stimulus location across temperatures. Using a linear decoder with leave-one-out cross-validation (see Methods), we found that decoding accuracy was high and unaffected by temperature (Figure 4G). Overall, despite a reported decrease in tectal responses above 33°*C* ^40^, within the ecological range tectal responses to prey-like stimuli and decoding performance were stable across temperatures. This robustness parallels the temperature invariance observed in hunting performance, as robust tectal responses drive the visuomotor transformations required for hunting behavior ^29,31,37,38,39^.

### Neural activity temporally scales with temperature

Given the temperature-dependent temporal scaling of hunting behavior, we asked whether the temporal dynamics of neural activity showed a similar pattern. We first examined the evoked sensory response dynamics across all tuned neurons at each temperature. As temperature increased, peak response emerged earlier (Figure 5A, Figure S5A). To quantify this effect, we measured the response time to peak, defined as the time from stimulus onset to the peak of the average neural response across neurons, at each temperature. Response time to peak was shortest in the hot condition and longest in the cold condition (Figure 5B). These changes in response time to peak were not due to differences in peak response amplitude, which was similar between hot and intermediate conditions and only slightly lower in the cold condition (Figure 5C). These results indicate that, at the population level, sensory tectal responses scale temporally with temperature, paralleling the behavioral effect.

**Figure 5:**
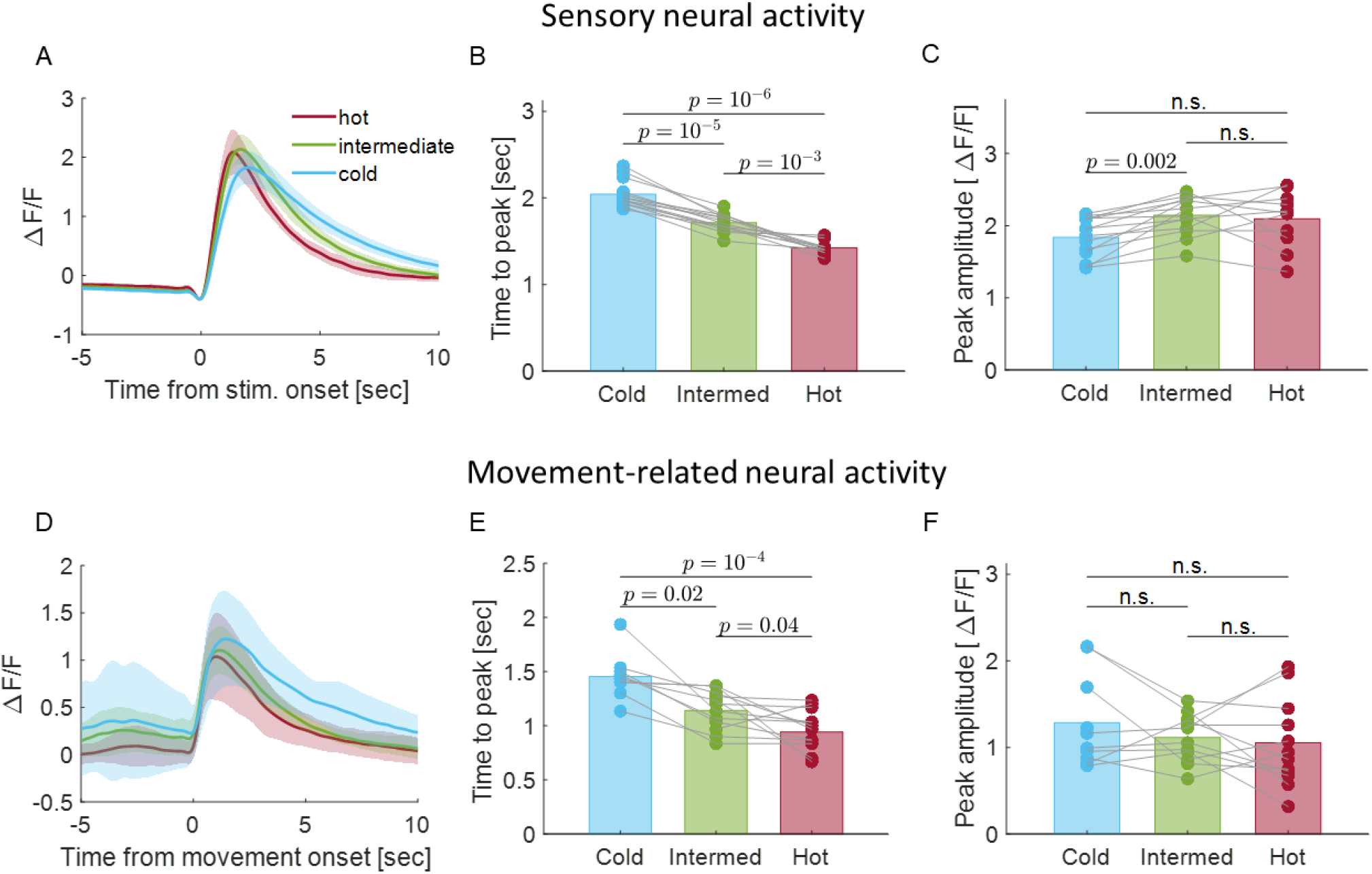
Temperature modulates sensory and motor neural response time. **(A)** Mean sensory response of tuned neurons to stationary visual stimuli across fish. Peak response appeared earlier in hot temperature and was progressively delayed as temperature decreased. Time zero is the stimulus onset. Shaded areas indicate the standard deviation across fish. **(B)** Response time to peak decreased with increasing temperature across fish. **(C)** Peak response amplitude was lower in the cold condition compared to the intermediate condition, but not different between intermediate and hot conditions. **(D)** Mean movement-locked neural activity of hindbrain neurons across fish. Peak response appeared earlier in hot condition and was delayed as temperature decreased. **(E)** Response time to peak decreased with increasing temperature across fish. **(F)** Peak response amplitude did not differ across conditions.

We next asked whether this temporal scaling extended beyond sensory responses to motor-related activity. During the experiments, fish exhibited spontaneous tail movements, and we collected neural activity locked to these movements. To identify the neurons whose activity was correlated with tail movement onset, we used a movement-locked regressor (see Methods, Figure S5B). These neurons were predominantly located in the hindbrain and comprised on average 48.3% of the detected neurons in this brain area (554±40 neurons, mean±SEM). As with sensory responses, the temporal dynamics of these motor-related responses varied with temperature (Figure 5D, see Methods). Time to peak response was again shortest in the hot condition and longest in the cold (Figure 5E), while response amplitude, which highly varied compared to the sensory responses, did not differ across temperatures (Figure 5F). Thus, both sensory and motor-related responses exhibit temperature-dependent temporal scaling.

Temporal scaling of neural responses was also evident at the single-neuron level. To examine this, we tracked individual neurons recorded at all three temperatures by acquiring a 300*μm* structural volume after functional imaging and registering each plane to this reference stack (see Methods). This allowed us to track the same neurons recorded across temperatures and examine their temporal dynamics (Figures S5C,D). Among these matched neurons, response time to peak decreased with temperature (Figure S5E), while response amplitude did not differ across temperatures (Figure S5F). Together, temperature-dependent temporal scaling is preserved not only at the population level but also at the level of individual neurons.

Does temperature affect sensory and movement-related responses in a similar manner? Since response time estimates were anchored to different reference points (stimulus onset for sensory responses and movement onset for motor responses), direct comparisons of their temporal dynamics were confounded. To overcome this, we assessed the temporal dynamics of neural activity using the autocorrelation of each neuron, which does not rely on external anchors (see Methods). While temporal dynamics scaled similarly with temperature in both populations (Figures S5G,H; two-way ANOVA interaction: *p* = 0.38), temporal dynamics were slower in the motor-related responses compared to the sensory responses (paired t-test, Benjamini-Hochberg correction, hot: *p* = 0.021; intermediate: *p* = 10^−4^; cold: *p* = 0.005). These findings suggest that temperature may exert a consistent global effect on neural dynamics, while preserving differences between sensory and motor-related populations.

Could these temporal differences reflect temperature-dependent changes in calcium indicator dynamics? Although a modest 10 ms change in indicator rise time has been reported over a 12°*C* range (from 25°*C* to 37°*C*) ^41^, we observed a 200 ms difference in neural response timing across a much smaller 4°*C* range (from 27.5°*C* to 31.5°*C*). This suggests that indicator kinetics alone cannot account for the effect. To further support it, we compared the temporal dynamics of neural responses to stationary and moving stimuli by fitting the rising phase of the calcium transient with an exponential function and extracting the rise time parameter. Across all temperature conditions, moving stimuli evoked faster responses than stationary stimuli (Figure S5I). This suggests that the indicator was capable of capturing meaningful stimulus-dependent differences in neural dynamics across temperatures and was not a limiting factor in our measurements. Overall, both sensory and motor-related neural responses exhibit temperature-dependent temporal scaling, with faster responses in the hot condition and slower responses in the cold, paralleling the temporal scaling observed in hunting behavior.

### A neural temporal parameter can explain robust hunting behavior across temperatures

Our neural results show that neural activity accelerates with increasing temperature, as indicated by shorter response times (RT). In parallel, our behavioral results demonstrate that hunting larval zebrafish maintain a stable desired bout distance across temperatures by increasing tail beat frequency (freq) and shortening bout duration as temperature increases (Figure 6Ai). We hypothesized that this behavioral stability during hunt does not require active compensation, i.e., explicit adjustment of tail beat frequency and bout duration, but instead reflects an intrinsic, self-organizing property of the neural circuit. We illustrate this using a simple rate model ^42^. Given that oscillatory neural activity underlies tail movements ^43,44,45^, we modeled an oscillatory system in which the input represents a desired bout distance and the output drives tail oscillations. We asked whether temperature-dependent changes in a single neural parameter, such as the neural response time, could account for the observed coordinated modulation in oscillation frequency and duration of hunting movements, ultimately preserving a consistent bout distance across temperatures.

**Figure 6:**
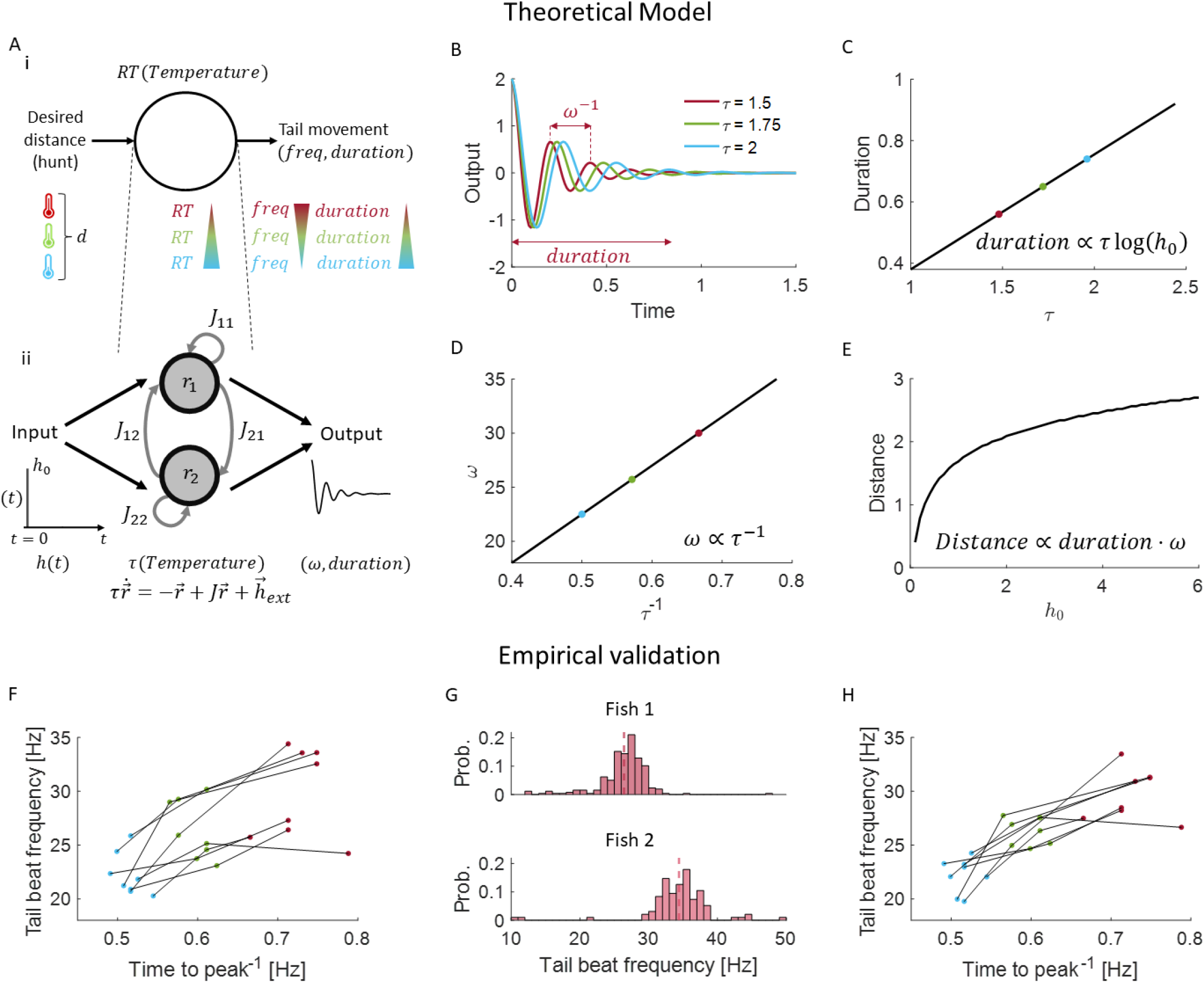
A single neural temporal parameter can account for the coordinated modulation of tail beat frequency and bout duration. **(A) i**. A schematic summary of the experimental results during hunting behavior. The distance (*d*) remained consistent across temperatures, while movement tail beat frequency (*freq*), movement duration, and response time of neural activity (*RT*) were modulated with temperature. **ii**. A simple rate model of two connected neurons with a connectivity matrix *J*, receiving an instantaneous input *h*(*t*), representing the desired distance, and producing an oscillatory output, driving tail oscillation. **(B)** Three example oscillatory outputs of the model for a fixed external input (*h*_0_) and three values of *τ*, illustrating that *τ* determines both the oscillation frequency and duration. As *τ* decreases, oscillation duration shortens and frequency increases, as indicated by cycle time (*ω*^−1^). Oscillation duration was defined as the time required for the amplitude to decay below a fixed threshold. All values are in arbitrary units. **(C)** For a given input, oscillation duration is linearly correlated with *τ*. Colored dots correspond to durations of oscillations generated by the model for the three *τ* values shown in (B). **(D)** Oscillation frequency (*ω*) was linearly correlated with *τ*^−1^and independent of the input. Colored dots correspond to oscillations generated by the model for the three *τ* values shown in (B). **(E)** The effective ‘distance’ generated by the oscillatory output, predominantly by the product of its duration and frequency, increases with the strength of the input *h*_0_, and is unaffected by *τ*. **(F)** Tail beat frequency as a function of neural response time to peak, measured from sensory responses, showing a similar trend in each fish (ANCOVA, non-significant interaction term *p* = 0.74). Each dot represents a single fish recording, and temperature conditions are indicated by colors. Black lines connect the same fish across temperature conditions. Fish that had both behavioral and neural recordings across all three conditions were included. **(G)** Example distributions of tail beat frequency from two head-fixed, tail-free fish under the hot temperature condition, illustrating inter-individual variability in baseline tail beat frequency. Dashed colored lines indicate the means. **(H)** Adjusted tail beat frequency linearly correlates with neural response time to peak, using a linear mixed model (LMM) (*R*^2^ = 0.74). This analysis empirically validates the model-predicted linear relationship between *τ*^−1^and *ω* as shown in (D).

The model consists of two interconnected neurons by a connectivity matrix (*J*) and a time constant *τ* that determines their response time. An instantaneous external input (*h*(*t*) = [*h*_0_, *t* = 0; 0, *t* > 0]), representing the desired movement distance, is applied to both neurons, and triggers oscillatory activity characterized by frequency (*ω*) and duration (Figure 6Aii). In this model, both oscillation duration and frequency are governed by *τ*, while duration additionally depends on the input strength, *h*_0_ (Figure 6B, see Methods). For a given input, oscillation duration scaled linearly with *τ* (Figure 6C), while oscillation frequency was inversely proportional to *τ* and independent of input (Figure 6D). The effective ‘distance’, computed as the product of oscillation duration and frequency (a proxy to movement velocity ^34,35,36^), remained invariant to *τ* and scaled with input strength (Figure 6E). Thus, as *τ* decreases, the circuit naturally produces shorter oscillation durations and higher frequencies, even when the input remains constant, mirroring the experimental results of hunting behavior. While this type of scaling is a general property of dynamical systems with a single time constant, the model provides a minimal framework for understanding how temperature-dependent neural dynamics can drive behavioral temporal scaling while maintaining spatial stability.

We next asked whether the observed changes in neural response time could linearly account for the modulation of tail beat frequency, as predicted by the model (Figure 6D). To directly link neural and behavioral dynamics, we analyzed simultaneously recorded neural activity and tail movements. As in freely swimming fish, head-fixed and tail-free fish showed shorter bout durations and higher tail beat frequencies with the increase in temperature (Figures S6A,B). Consistent with model predictions, tail beat frequency was linearly correlated with the neural response time, as extracted from neural activity (Figure 6F). While all fish showed a temperature-dependent increase in frequency (Figure 6F), baseline frequencies varied across individuals (Figure 6G, Figure S6C). A linear mixed-effects model, accounting for this inter-individual variability, confirmed a significant correlation between response time and tail beat frequency across fish (Figure 6H). These results support the prediction of the model: faster neural dynamics at higher temperatures directly drive higher tail beat frequencies and shorter bouts. Although response time is only a proxy for the true biophysical neuronal time constant, it still shows a linear relation with tail beat frequency. Crucially, this suggests that robust hunting behavior can emerge as a self-organizing property of temperature-dependent neural dynamics.

## Discussion

Animals are often required to maintain stable performance in critical behaviors despite environmental fluctuations. We show that larval zebrafish maintain robust hunting performance across a 10°*C* ecological temperature range. During hunting, behavioral dynamics exhibit substantial temporal scaling with temperature, yet key movement spatial parameters, such as bout distance and turn angles, remain remarkably stable. This stability is achieved through coordinated adjustments in tail dynamics, characterized by increased tail beat frequency and reduced bout duration as temperature increases. The temporal scaling observed in the behavior was extended to the neural activity of the population and single neurons, with neural activity accelerating in both sensory and motor domains as temperature increased. We suggested that the coordinated modulation of tail beat frequency and movement duration, which is required to maintain robust distance across temperatures, is governed by the observed neural temporal scaling. To illustrate it, we constructed a simple rate model showing that temperature-dependent changes in neural response time alone can account for the observed increase in frequency and decrease in duration, preserving effective movement distance without requiring active compensation. The model predictions were well fitted to the empirical data. These findings reveal a potential self-organizing mechanism through which temperature-sensitive neural circuits preserve the performance of critical behaviors, offering a general principle for behavioral robustness in dynamic environments.

Temperature influenced the system across multiple levels, from whole-animal behavior to the activity of individual neurons. In hunting behavior, temperature-dependent temporal scaling was evident at all timescales, from individual movement durations and IBIs to complete behavioral sequences. This scaling was mirrored in tail dynamics: as temperature increased, tail beat frequency increased and movement durations shortened across behavioral contexts (hunting, exploration, and in head-fixed condition) (Figure 3I, Figure S3D, Figure S6B). Similar temporal scaling was observed in neural activity, both at the population and single-neuron levels, consistent with well-established effects of temperature on membrane biophysics, ion channel kinetics, and synaptic transmission ^10,46,47,48,49^. These findings highlight the extensive impact of temperature and the importance of precise thermal control in experiments probing neural activity, behavior, or their interaction.

We implemented a simple rate model that generates oscillatory activity and showed that altering the neural time constant alone was sufficient to modulate both oscillation frequency and duration. This coordination preserves a consistent movement distance without requiring additional control during hunting. While minimal and one of many possible models, this model illustrates a general principle that a single temporal parameter can shape multiple kinematic features through intrinsic circuit dynamics. Importantly, the model also captures the temporal scaling observed during exploration. As in hunting, tail beat frequency increased with temperature during exploration (Figures S3B,E), and an inverse relationship between frequency and bout duration was observed (Figures S3C,F). While we did not rule out the possibility of active compensation, such as modulation by temperature-sensing neurons in the trigeminal ganglion ^50^, our model, nonetheless, offers a parsimonious and biologically plausible explanation for how behavioral robustness across temperatures could arise from intrinsic neural dynamics governed by a single parameter.

To estimate distance, the model uses the product of tail beat frequency and oscillation duration, based on prior evidence that tail beat frequency correlates with swim velocity ^34,35,36^. Interestingly, the variance of this estimated distance was lower than that of the actual measured distance of the freely swimming fish (Figure S6D), suggesting that factors beyond tail beat frequency and movement duration determine the actual movement distance. While a more detailed tail dynamics model (and potentially fin movements) is necessary to accurately extract movement distance, the current approach supports a key insight: despite simplifications, the model reveals that during critical behavior, such as hunt, zebrafish can maintain a desired travel distance by intrinsically adjusting both tail beat frequency and bout duration. This highlights a flexible yet robust possible mechanism for achieving behavioral stability across environmental conditions.

One possible contributor to the observed increase in tail beat frequency is the temperature-dependent change in water viscosity. From 32°*C* to 22°*C* water viscosity is reduced by approximately 20% ^51^. Previous work has shown that a much larger, 3.3-fold reduction in viscosity can increase tail beat frequency by only 2–4 Hz in zebrafish ^52^. In contrast, we observed an increase of over 12 Hz across our temperature conditions, corresponding to a relatively modest change in viscosity. This discrepancy suggests that while temperature-induced changes in viscosity may contribute to increased tail beat frequency, they are unlikely to fully explain the magnitude of the effect. Instead, our findings point to a dominant role for temperature-dependent modulation of neural activity in driving the observed changes in motor output.

Is temporal scaling in neural activity similar across all neurons? We found no significant interaction between temperature-dependent temporal response profiles and neuronal population type, suggesting that temperature similarly affects sensory and motor-related neurons. However, their temporal dynamics, as estimated from neuronal autocorrelation, differed (Figures S5G,H). Specifically, motor-related neurons showed slower dynamics, potentially reflecting longer response delays. These delays may arise from synaptic transmission, accumulated sensory processing time, or heterogeneous functional roles within the motor population, contrasting with the more stereotyped, evoked sensory responses. More broadly, our response time metric serves as an indirect proxy for intrinsic neural dynamics and does not directly capture biophysical time constants; further research is required to determine whether this form of temporal scaling is consistent across all neuron types or reveals important functional differences. Whether a common, brain-wide temperature-sensitive time constant shapes activity, or whether scaling is confined to specific oscillation-generating circuits, the consistency of tail beat frequency within individual fish suggests that network-level synchrony is preserved across temperatures, at least in the circuits driving these behaviors.

Our findings reveal a fundamental distinction in how zebrafish adapt their behavior to temperature across contexts. During exploration, bout distance increases with temperature, whereas during hunting, spatial precision is maintained, with bout distances remaining stable. This raises two key questions: what is the functional significance of this context-dependent modulation, and what mechanism underlies it? A likely functional explanation is that the stability of hunting reflects a fixed goal, prey capture, that remains consistent across temperatures, requiring precision regardless of thermal conditions. In contrast, exploratory behavior, lacking a specific external target, may be guided by shifting internal goals between temperature conditions. For instance, at higher temperatures, fish might increase movement to seek cooler areas, while at lower temperatures, they may conserve energy by swimming less^4^. Thus, temperature-dependent changes in exploratory behavior may reflect goal flexibility and illustrate context-dependent modulation of selected actions.

What mechanism could account for this context-dependent modulation? Specifically, how can zebrafish vary traveled distance with temperature during exploration but not during hunting? Although both tail beat frequency and bout duration vary significantly with temperature (two-way ANOVA, temperature effect, *p* = 0 for both), only movement duration differs across behavioral contexts (two-way ANOVA, context effect, *p* = 0) while tail beat frequency remains similar across contexts (two-way ANOVA, *p* = 0.76). This suggests that bout duration is modulated differently during exploration to achieve different distances across temperatures. Two lines of evidence from our data support a potential mechanism. First, the product of tail beat frequency and duration, an estimate of bout distance, shows significantly lower variance during hunting than exploration (F-test, *p* = 0.002), indicating tighter distance regulation. Second, fitting the relationship between tail beat frequency and bout duration with a hyperbola (assuming a constant product, which approximates a fixed distance), explains more variance during hunting than exploration (Figure S6E, hunting: R^2^ = 0.74; exploration: R^2^ = 0.58). This suggests that during hunting, movement duration and frequency are more tightly coordinated to preserve a consistent movement distance, supporting the idea of context-dependent motor control precision. Although our neural recordings did not allow accurate inference of behavioral context, future studies that combine neural activity with extended behavioral tracking, including eye movements, could help uncover the mechanisms of context-dependent action selection. In this framework, temperature serves not only as a naturalistic perturbation but also as a powerful tool to reveal how internal states modulate sensorimotor transformations. Our findings, in this context, lay the groundwork for uncovering the neural basis of context-dependent action selection.

## Acknowledgements

We thank Harold Burgess, Hagar Lavian, Inbal Goshen, and Mati Joshua for their valuable feedback on earlier versions of the article. We are also grateful to Michael London, Mark Shein-Idelson, and the Avitan lab members for inspiring and stimulating discussions. Finally, we thank the fabrication laboratory team at ELSC for their assistance in refining our experimental setup.

## Methods

### Zebrafish maintenance

Zebrafish larvae (Danio rerio) expressing elavl3:H2B-GCaMP6s^53^ were collected and raised according to established procedures ^54^. Larvae were maintained at 28°*C* under a light and dark cycle of 14/10 hours and were fed live rotifers (Brachionus plicatilis) daily, from 5 dpf. Behavioral experiments were conducted at 13 dpf, and calcium imaging experiments were performed at 14 dpf. At this age, the sex of the fish cannot be determined. All procedures were performed with approval from The Hebrew University of Jerusalem Animal Ethics Committee. The Hebrew University is accredited by the Association for Assessment and Accreditation of Laboratory Animal Care International (AAALAC).

### Experimental feeding assay

Individual larvae (13 dpf, n=30, 10 per temperature condition) were placed in a transparent-bottom plate (20 mm diameter and 2.5 mm depth, CoverWell Imaging Chambers, Catalogue number 635031, Grace Biolabs) with 30 Paramecia for 15 minutes. The plate was illuminated from above with visible white light and from below with infrared LEDs beneath the plate (L850 nm, LDR2-100IR2-850-LA, powered by PD3-3024-3-PI, CCS Inc., Kyoto, Japan). Natural larval behavior was recorded from above, using a high-speed camera (MIkrotoron 679 4CXP) at 500 frames per second and 45 microns per pixel. Room temperature was adjusted for each temperature condition (16°*C*, 25°*C* and 29°*C*), and water temperature was precisely maintained using a thermo-plate (Tokai Hit) to achieve target temperatures (cold (21.96±0.68°*C*), intermediate (26.98±0.54°*C*), and hot (32.08±0.88°*C*)). Larvae were habituated to the experimental temperature for at least 20 minutes in a separate dish prior to behavioral imaging. Water temperature was recorded immediately before and after each recording session.

### Behavioral tracking and feature extraction

Fish behavior was tracked using a custom-developed algorithm ^23^. The algorithm extracted key features of the fish posture and movement, including the fish contour, the tail midline (from swim bladder to tail tip), eyes postures, as well as prey location. To identify bouts of movements, we selected 15 equidistant points from the tail tip along the fish midline (out of 100, spanning the tail tip to swim bladder) and calculated their mean velocity based on changes in the Euclidean distance between consecutive frames. A velocity threshold was applied to detect candidate movement frames. Potential detected bouts defined by fluctuations in velocity were validated using the velocity standard deviation over time (Figure S1B). Bouts boundaries were refined using singular value decomposition (SVD) of the tail posture, identifying the precise start and end frames based on the squared sum of the first two components. Detected bouts shorter than 16 frames (32 ms) were excluded. A total of 5818 bouts were detected and analyzed during hunt (1686, 2413, 1719 in the hot, intermediate and cold conditions, respectively). Hunting events were segmented using eye convergence and divergence to detect hunt onset and offset ^23,30^, and our automated segmentation was manually validated. For exploration segments, 2-second windows were selected between hunting events, with the number of windows matching to the number of detected hunting events. In these segments, there was no eye convergence or interaction with the prey. During exploration, 4465 bouts were detected and analyzed (1692, 1794, 979 in the hot, intermediate and cold conditions, respectively). For the estimation of mean prey velocity (Figure S2C), the last bout of the event was excluded as the prey could only be partially detected during these bouts. For analyses of the interaction between the fish and its prey, only successful hunting events were included since unsuccessful events show a less consistent relationship between the fish and its prey ^32^.

### Hunting events with matched Paramecia velocity across temperatures

To determine whether the changes in bout and IBI durations were driven by temperature-dependent prey velocity, we analyzed Paramecia movement across temperature conditions. Paramecia moved more slowly in the cold condition compared to the intermediate and hot conditions, although velocity distributions overlapped substantially (Figure S2C). To isolate temperature effects on fish behavior, we selected hunting events in which Paramecia velocity fell within a shared range, spanning the mean velocities of cold and intermediate conditions, and was not statistically different across temperature conditions. We then calculated fish movement statistics within this subset of events (Figures S2D-I).

### Tail beat frequency estimation

To evaluate tail beat frequency, we first quantified tail postures within each bout and subsequently assessed oscillation within these postures. More specifically, for each fish, we extracted the positions of 100 equidistant points along the tail midline during every detected bout and calculated the angles between each segment with a reference axis, forming a matrix of tail postures across all bouts. Singular value decomposition (SVD) was then applied to this matrix to identify the principal components, representing the dominant modes of tail posture during movements. For each bout, we applied a Fourier transform to the time series of the first two principal component coefficients, using a Tukey window to minimize edge artifacts (with a Tukey parameter of 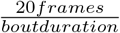). We then identified the dominant tail frequency as the peak frequency within the 10-50Hz range. This approach captured the most prominent oscillatory component of tail movement during each bout.

### Simultaneous two-photon calcium imaging and tail movements recording

A single 14 dpf larva was embedded in 2.5% low-melting agarose at the center of a 33 mm diameter dish with the tail free to move (n=14). Neural activity was recorded using single-plane two-photon calcium imaging (Nikon objective, CFI ×16, 0.8 NA), capturing a 705*×*352*μm* field of view at a rate of 29.946 frames per second. Imaging was performed at a depth of 65−75*μm* below the skin surface, consistent with depths used in previous studies of evoked tectal responses ^29,55^. Simultaneous tail movements were recorded from below using an infrared camera (FLIR Grasshopper 3) at 500 fps, with illumination provided by 850nm infrared LEDs.

Visual stimuli were projected using a projector onto high contrast diffusion paper attached to the dish using Optoma ML750ST with 600 nm long-pass filter. Water temperature was controlled via a thermoplate (Tokai Hit) and monitored using a thermocouple inserted into the plate. Each fish was recorded under all three temperature conditions (cold (23.28±0.74°*C*), intermediate (27.52±0.26°*C*), and hot (31.62±0.41°*C*), starting in either the cold (n=7) or the hot (n=7) condition. Temperature conditions under the microscope varied on average across conditions by 0.75°*C* from those of the behavioral experiments. Recordings were spaced by one hour of habituation per condition. Each session began with a 5-minute dark phase, followed by visual stimuli presented in pseudo-random order.

Stationary stimuli were presented in four angular positions (−60°,−30°,−30°,−60° relative to fish midline), while moving stimuli were shown in two directions (from −55° to−10° and from 55° to 10° at 30° per second (n=8) or 90° per second (n=6)). Each stationary stimulus was presented 20 times and each moving stimuli 15 times, for a total of 80 stationary stimuli and 30 moving stimuli. Stimuli were separated by 20-second intervals, with 35-second pauses between blocks of 22 stimuli. Three hot condition neural recordings were omitted from the dataset due to instability of the sample (one as the first recordings and two as the last). Behavioral recordings of a single fish across all temperature conditions and two additional recordings in the cold condition were omitted from the analysis due to low signal-to-noise ratio (SNR) of behavioral imaging.

For two of the two-photon recorded fish, neural structural scans of the entire brain were acquired following functional imaging (300 imaging planes, 1*μm* apart, 1 second per plane). Functional recordings of these fish were registered to the structural volume, aligning all detected neurons within a common coordinate system. Registration was performed using Advanced Normalization Tools (ANTs^56^). This revealed that depth differences across three temperature recordings for each fish were less than 2*μm*. Using this shared coordinate system, we were able to identify individual neurons that were active across all temperature conditions (Figure S5D).

### Neural recordings preprocessing

Raw two-photon calcium imaging data were processed using Suite2p ^57^, which included motion correction and cell detection. Δ*F*/*F* traces were computed for each cell, with the baseline defined as the 8th percentile of a moving window over 5000 frames (167 seconds). Traces were then smoothed using a zero-phase low-pass filter (filtfilt Matlab function) and z-scored. For each recording, the midbrain and hindbrain were manually annotated. Behavioral data, recorded during the two-photon neural imaging, showed spontaneous tail movements at rates of 3.8, 2.9, 2.1 movements per minute in hot, intermediate and cold conditions, respectively. These movements were not correlated with the stimuli (*p* = 0.80). Behavioral recordings were binarized, and tail features were extracted using a modified version of the custom-made tracker developed for behavioral recording of freely moving fish. To minimize the potential influence of neural activity originating from preceding or subsequent movements, particularly for response time estimates of motor-related activity, we focused on isolated bouts, which were separated by at least 3 seconds from adjacent bouts. A total of 4919 bouts were detected and analyzed (2224, 1658, and 1037 in the hot, intermediate, and cold conditions, respectively). Of these, 889 bouts were separated by at least 3 seconds from adjacent bouts and were included in the neural analysis (307, 305, and 277 in the hot, intermediate, and cold conditions, respectively).

### Forward bouts estimation

To estimate forward bouts, we computed a laterality index defined as the average angle between the tail tip and the swim bladder during the initial 70ms of each bout ^58,59,60^. This index was found to be correlated with turn angle in freely swimming fish and has been previously used for the classification of forward versus turn bouts in head-fixed fish. Subsequently, for each fish, we calculated the distribution of the absolute laterality index and fitted it with a dual Gaussian distribution, taking the intercept between the Gaussians as the threshold for classification between forward and turn bouts.

### Tuned neurons identification using regressors

To identify tectal neurons whose activity was correlated with presented stationary stimuli, we constructed regressors for each unique angular position and calculated Pearson’s correlation values for each neuron with each regressor. Regressors were generated by convolving stimulus onset times with an exponential decay kernel, designed to match the dynamics of GCaMP6s. Preliminary analysis revealed temperature-dependent differences in response dynamics, specifically in response time to peak and decay rates, prompting us to refine the regressors by incorporating a stimulus-onset delay (1.0, 1.27, and 1.5 seconds for hot, intermediate, and cold conditions, respectively) and adjusting the decay constants of the exponential kernel (1.2, 1.5, and 1.9 seconds, respectively). To determine statistical significance, we generated a null distribution of correlation values by randomly shuffling stimulus labels 100 times. For each shuffle, new regressors were created, and correlations were recomputed. Neurons were considered stimulus-correlated if their correlation exceeded the 97th percentile of the null distribution.

Movement correlated neurons were identified using a similar approach, based on correlation with regressors constructed by a convolving movement onsets and a box kernel of 3 seconds duration. Box kernel was chosen because the onset of movement-related neuron activity in the hindbrain was less temporally characteristic compared to tectal responses to visual stimuli. To determine a correlation threshold, we generated a null distribution using regressors with the same number of movement events and the same kernel, but with randomly shuffled onset times. Neurons were considered movement-correlated if their correlation exceeded the 99th percentile of the null distribution. To assess the temporal dynamics of movement-related neural activity, we focused on isolated movements spaced at least 3 seconds apart, minimizing potential influence of neural activity driven from preceding or subsequent movements.

### Decoding stimulus angular position from neural activity

To decode the angular position of stationary stimuli from neural activity, we used a linear discriminant analysis (LDA) decoder. For each fish, we computed the mean activity of all identified stimulus-tuned neurons within a 3-second window following each stimulus presentation, generating a population activity matrix. The LDA decoder was trained using a leave-one-out cross-validation approach, iterating over all stimulus presentations. Decoder performance was quantified as the proportion of correctly predicted stimulus positions.

### Calculating neuronal trace autocorrelation to compare temporal dynamics

To assess neuronal temporal dynamics of sensory responses and motor-related neural responses, without relying on different temporal anchors such as stimulus or movement onsets, we calculated the autocorrelation of each neuronal activity trace (Δ*F*/*F*) and fitted the initial 7 seconds of the autocorrelogram with a decaying exponent (Matlab fit, 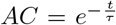). The mean autocorrelation decay factor across all neurons in each population was computed per fish (Figures S5G,H).

### A simple rate model for generating oscillatory activity

The model consisted of two neurons (whose firing rates are *r*_1_ and *r*_2_). The rate of each neuron was determined by the following dynamical equation ^42^:

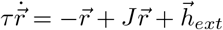

Where *τ* is the time constant of the neuron, 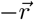 is the leak term, *J* is the connectivity matrix of the neurons, and 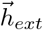 is the input to the system. We can simplify the equation by defining *M* = − *I* + *J* (where *I* is the unity matrix) to get:

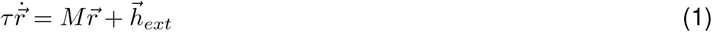

We assume *M* is a 2×2 real matrix with complex eigenvalues. For this case, the eigenvectors of M can be written as:

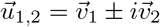

with corresponding eigenvalues:

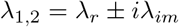

We assume that the system is stable, i.e., the dynamics do not diverge, giving us λ_*r*_ > 0.

We decomposed the vectors as a sum of the eigenvectors of M:

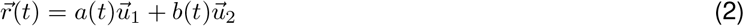

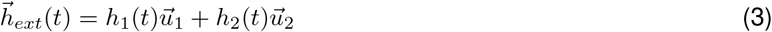

For simplicity, we defined the input to both neurons to be equal, *h*_1_(*t*) = *h*_2_(*t*) = *h*(*t*). The same qualitative results can be achieved without this simplification. Thus, the input to the system is:

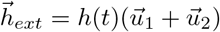

We consider a transient input of magnitude *h*_0_ applied at time *t* = 0:

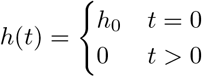

Plugging (2) and (3) in (1), yielded:

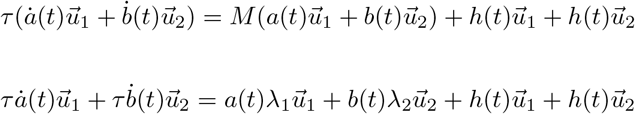

Multiplying by the left eigenvector corresponding to 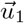 :

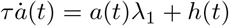

which has a unique solution:

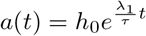

Multiplying by the left eigenvector corresponding to 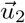 :

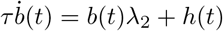

which has a unique solution:

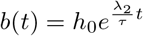

Plugging the results back to (2):

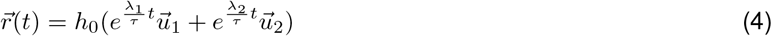

Since 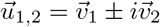, we can plug this into (4) to get:

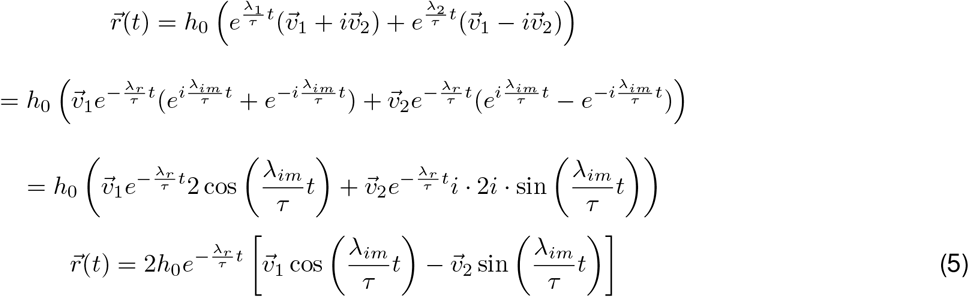

### Interpreting the solution

In equation (5), the expression outside the squared parentheses represents an overall amplitude that decays with time, and the expression inside the parentheses represents an oscillatory component with a typical frequency. Therefore, dynamics have a frequency which is inversely related to the neuronal time constant *τ* :

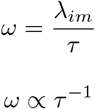

If we assume that tail dynamics are abolished when the firing rate is below a threshold *ϵ*, the decay time (or the duration of the oscillation) for the system is achieved when:

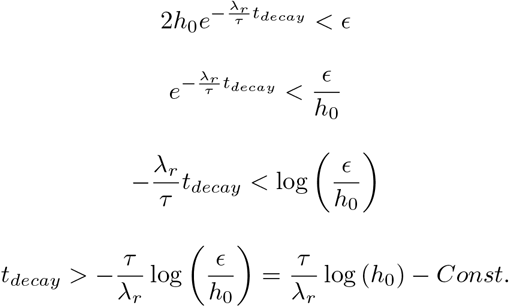

Therefore, the decay time (or the effective duration of the oscillation) is proportional to:

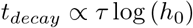

In conclusion, as *τ* increases (longer response time), oscillation duration increases and frequency decreases. The estimated distance from the model is:

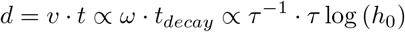

Meaning that the distance is proportional to the input:

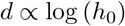

Therefore, the input to the system determines the distance, and is independent of the response time *τ*.

### Statistical analysis

To compare behavior across three temperature conditions, we used analysis of variance (ANOVA), accompanied by post-hoc multiple comparisons (Tukey-Cramer correction) for comparison between each pair of conditions. To compare neural activity across three temperature conditions, since the same fish was recorded in each condition, we used repeated-measures ANOVA. For this test, sphericity was tested using Mauchly’s Test of Sphericity, which was not significant in all cases. Following repeated-measures ANOVA, post-hoc multiple comparisons (Tukey-Cramer correction) were used to compare each pair of conditions. To determine the interaction between two behavioral contexts, such as hunting versus exploration, and temperature conditions, a multi-way ANOVA was used. The Kolmogorov–Smirnov test (KS-test) was applied to determine differences between distributions, with a Bonferroni correction for multiple comparisons. A permutation test (Figures 3D-F) was performed by shuffling the labels of the temperature condition across all data points 100000 times, each time calculating the Pearson correlation between Δ*θ* and *ϕ*_*prey*_ and extracting the maximal correlation difference between groups. P-value was then computed as the proportion of permuted differences that were greater than the observed differences.

