## Supplementary Information for "It’s about time: neural temporal scaling accounts for robust hunting behavior across temperatures"

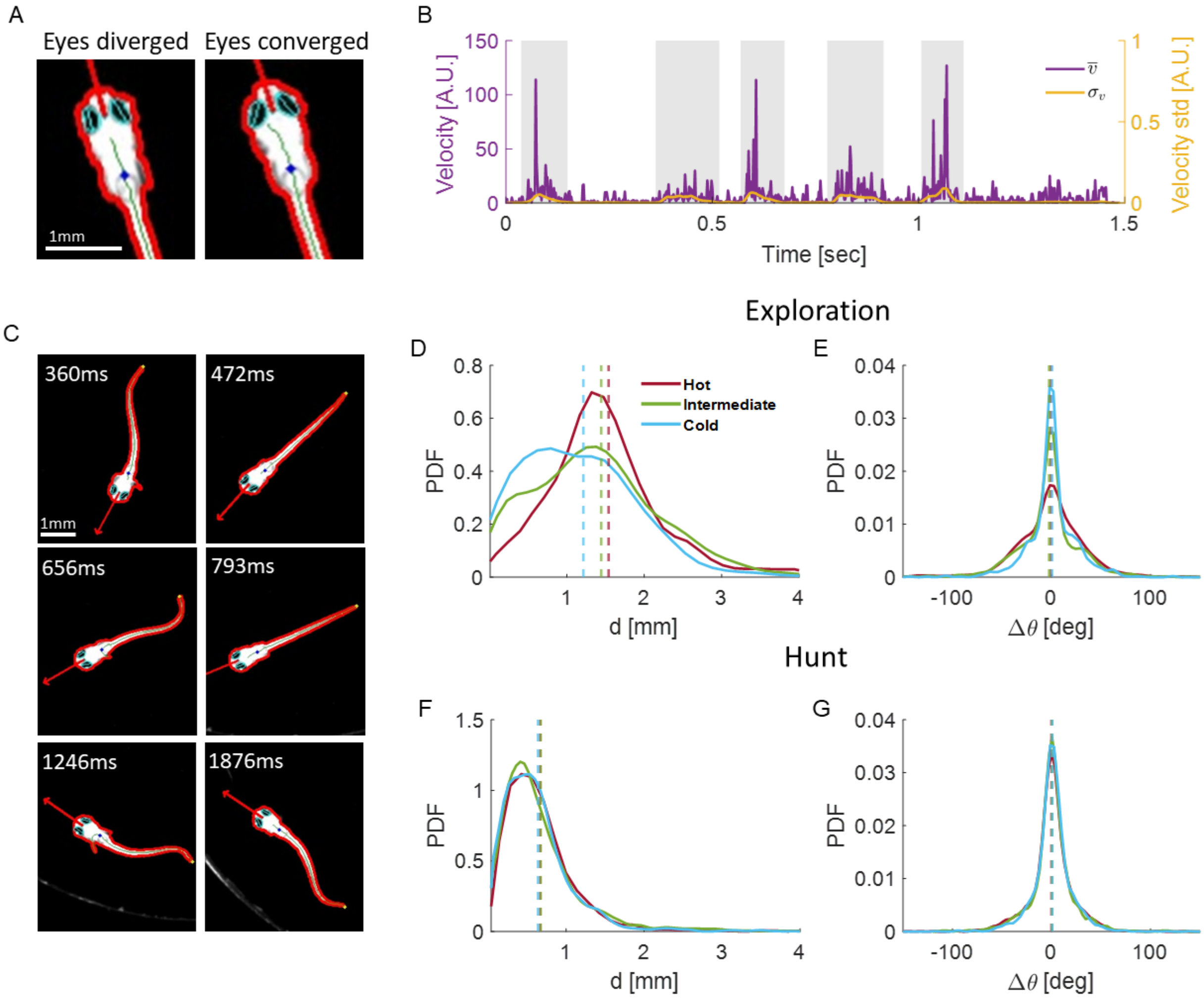


**Figure S1: Spatial movement statistics are modulated by temperature during exploration but remain robust during hunt. (A)** Fish eyes were tracked and fitted with ellipses (cyan), and the angle between their major axes was used to quantify eye convergence. Two example frames showing diverged eyes during exploration (left, $12.9°$) and converged eyes during hunt (right, $60.7°$). **(B)** Bout detection over an example hunting event. Detected bouts are marked in gray shaded areas. The mean velocity of 15 equidistant points out of 100 total points along the tail increased when the tail moved (purple trace). The standard deviation of this velocity across 15 points during the hunt (yellow trace) provided a smoother measure to segment movements. **(C)** Example frames during fish exploration, where eyes are diverged. **(D)** Bout distance distributions pooled across fish during exploration differed significantly across temperature conditions (Bonferroni-corrected KS-test, hot vs. intermediate $p = 10^{-8}$, intermediate vs. cold $p = 10^{-7}$, hot vs. cold $p = 10^{-25}$). Dashed vertical lines indicate the mean in each temperature condition. **(E)** Change in heading angle distributions pooled across fish during exploration differed significantly across temperatures (Bonferroni-corrected KS-test, hot vs. intermediate $p = 10^{-7}$, intermediate vs. cold $p = 10^{-4}$, hot vs. cold $p = 10^{-5}$). **(F)** Bout distance distributions pooled across fish during hunt were similar across temperatures (Bonferroni-corrected KS-test, hot vs. intermediate $p = 0.38$, intermediate vs. cold $p = 0.49$, hot vs. cold $p = 0.82$). **(G)** Change in heading angle distributions pooled across fish during hunt were similar across temperatures (Bonferroni-corrected KS-test, hot vs. intermediate $p = 1$, intermediate vs. cold $p = 1$, hot vs. cold $p = 0.48$).

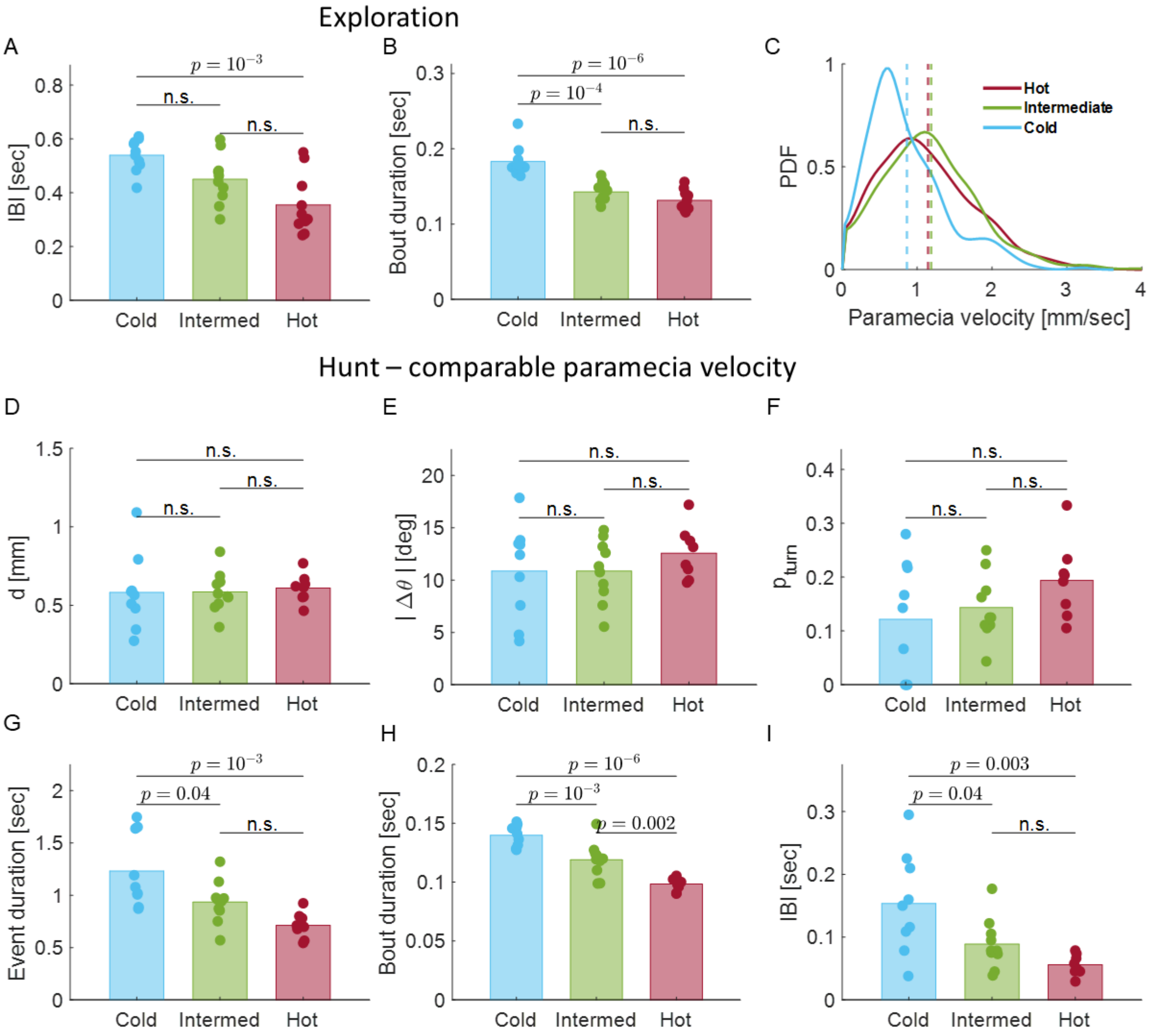


**Figure S2: Temperature-dependent modulation of movement temporal statistics is independent of paramecium velocity.** **(A)** During exploration between hunts, IBIs were shorter in the hot condition compared to the cold condition. **(B)** During exploration between hunts, bout duration was shorter in the intermediate and hot conditions compared to the cold condition. **(C)** Paramecia velocity distributions, pooled across recordings, differed significantly between cold and intermediate, and between cold and hot conditions, but were similar between hot and intermediate conditions (Bonferroni-corrected KS-test $p = 10^{-12}$, $p = 10^{-8}$, $p = 0.44$ respectively). Dashed vertical lines indicate the mean velocity for each temperature condition. Despite these differences, the distributions also overlapped substantially. To determine whether temperature-dependent changes in spatial movement statistics are driven by differences in Paramecia velocity, we analyzed a subset of events in which prey velocity fell within the overlapping range of the distributions, specifically between the mean values of the cold and intermediate conditions (0.78–1.31 mm/sec). For these events, prey velocity did not differ significantly across temperatures (ANOVA $p = 0.13$). **(D)-(F)** Fish movement spatial parameters, in events in which Paramecia velocity did not differ across temperatures, remained stable. (D) Bout distance ($d$) did not differ. (E) Absolute turn angles did not differ. (F) Turn probability did not differ. **(G–I)** Movement temporal parameters scaled with temperature even when paramecia velocity did not differ across temperatures. (G) Event duration was shorter in the hot compared to the cold condition. (H) Bout duration was shorter in the hot condition compared to the cold condition. (I) IBI duration was shorter in the hot condition compared to the cold condition.

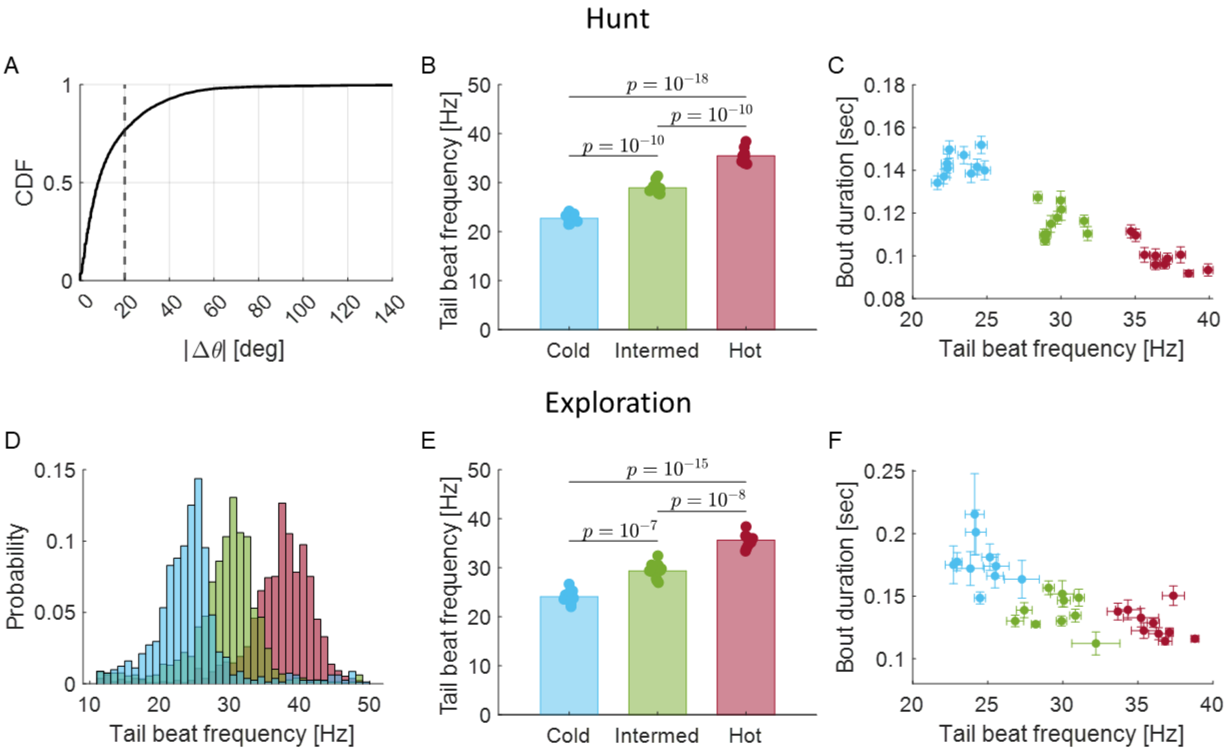


**Figure S3: Coordinated temperature-dependent modulations in tail beat frequency and bout duration. (A)** A cumulative distribution of the change in heading angle following a movement while hunting; most movements were forward movements. **(B)** Tail beat frequency significantly increased with temperature during hunt. **(C)** Tail beat frequency was inversely correlated with bout duration. Error bars are SEMs, Pearson correlation $\rho = -0.93$; $p = 10^{-12}$). **(D)** During exploration, tail beat frequency distributions differed significantly across temperatures (Bonferroni-corrected KS-test, hot vs. intermediate $p = 10^{-291}$, intermediate vs. cold $p = 10^{-162}$, hot vs. cold $p = 10^{-307}$). **(E)** During exploration, tail beat frequency significantly increased with temperature. **(F)** During exploration, bout duration was inversely correlated with tail beat frequency. Error bars are SEMs, Pearson correlation $\rho = -0.77; p = 10^{-6}$).

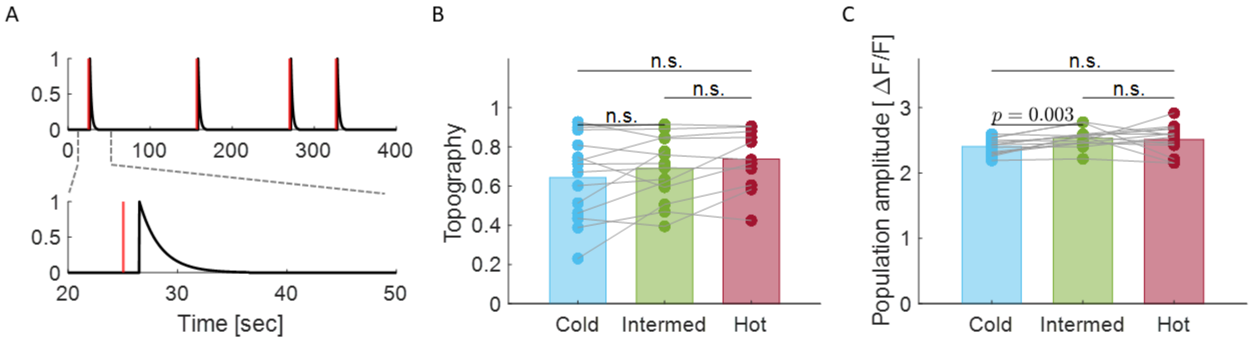


**Figure S4: Neural evoked sensory response is robust across temperatures. (A)** An example regressor used to identify neurons tuned to stationary visual stimuli. Red lines indicate stimuli onset. The regressor was constructed by convolving these stimuli onset with an exponential decay matching GCaMP6s dynamics. **(B)** Topography of sensory neural responses, measured as the correlation between neuronal tuning and position along the anterior-posterior tectal axis (see Methods), remained stable across temperature conditions. **(C)** Population response amplitude, calculated as the mean amplitude of recruited tectal neurons following stimulus presentation, was reduced in the cold condition compared to the intermediate condition, but was similar between hot and the other two conditions.

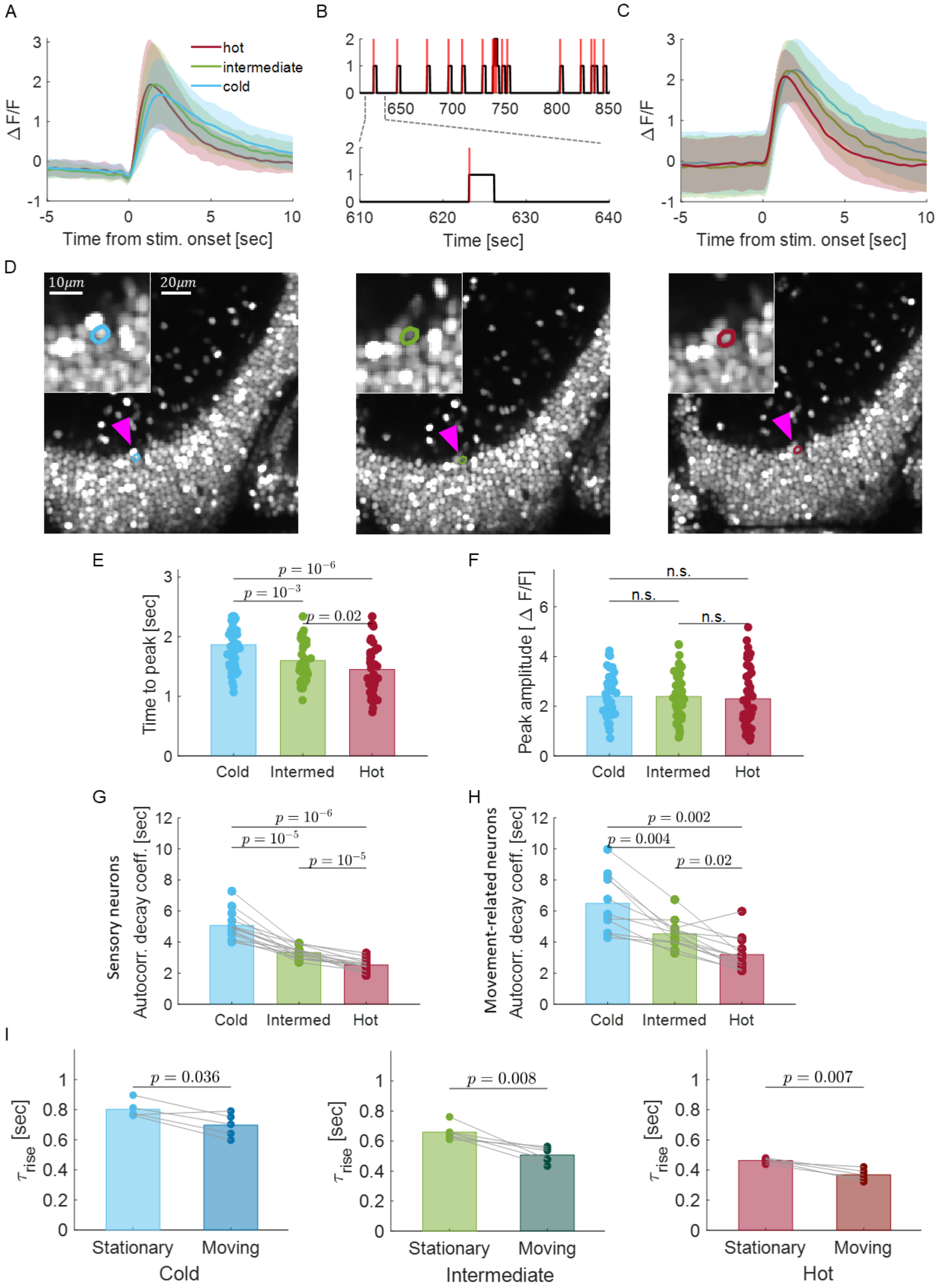


**Figure S5: See caption on next page.**

**Figure S5: Temperature-dependent neural temporal scaling at the population and single neurons level. (A)** Mean sensory response of tuned neurons to stationary visual stimuli of a single fish. Peak response appeared earlier in hot temperature and was progressively delayed as temperature decreased. Time zero is the stimulus onset. Shaded areas indicate standard deviations across neurons. **(B)** An example regressor used to identify neurons locked to the onset of movements. Red lines indicate movement onsets. The regressor was constructed by convolving these movement onsets with a 3-second rectangular ("box") function. **(C)** Mean sensory response across 45 tuned neurons that were detected and matched across temperature conditions in a single fish. The peak response emerged faster as the temperature increased. Shaded areas are standard deviations across neurons. **(D)** The location of an example neuron, detected across cold, intermediate, and hot conditions (right to left), indicated by a magenta arrowhead. **(E)** Response time to peak decreased with increasing temperature for single sensory neurons identified across all temperature conditions. Connecting lines were omitted for clarity due to the large number of neurons. **(F)** Peak response amplitude did not differ across conditions for single sensory neurons identified across all temperature conditions. **(G)** The exponential decay coefficient of neuronal autocorrelograms, calculated for sensory neurons, scales with temperature, reflecting temporal acceleration. **(H)** The exponential decay coefficient of neuronal autocorrelograms, calculated for each movement-related neuron, scales with temperature, reflecting temporal acceleration. **(I)** Fitted exponential rise time ($\tau_{rise}$) was shorter for moving stimuli ($90^{\circ}$ per second) than stationary stimuli across all temperatures, suggesting that the indicator faithfully tracks neural activity with sufficient temporal resolution to reflect varying stimulus speed (paired t-test, Benjamini-Hochberg correction).

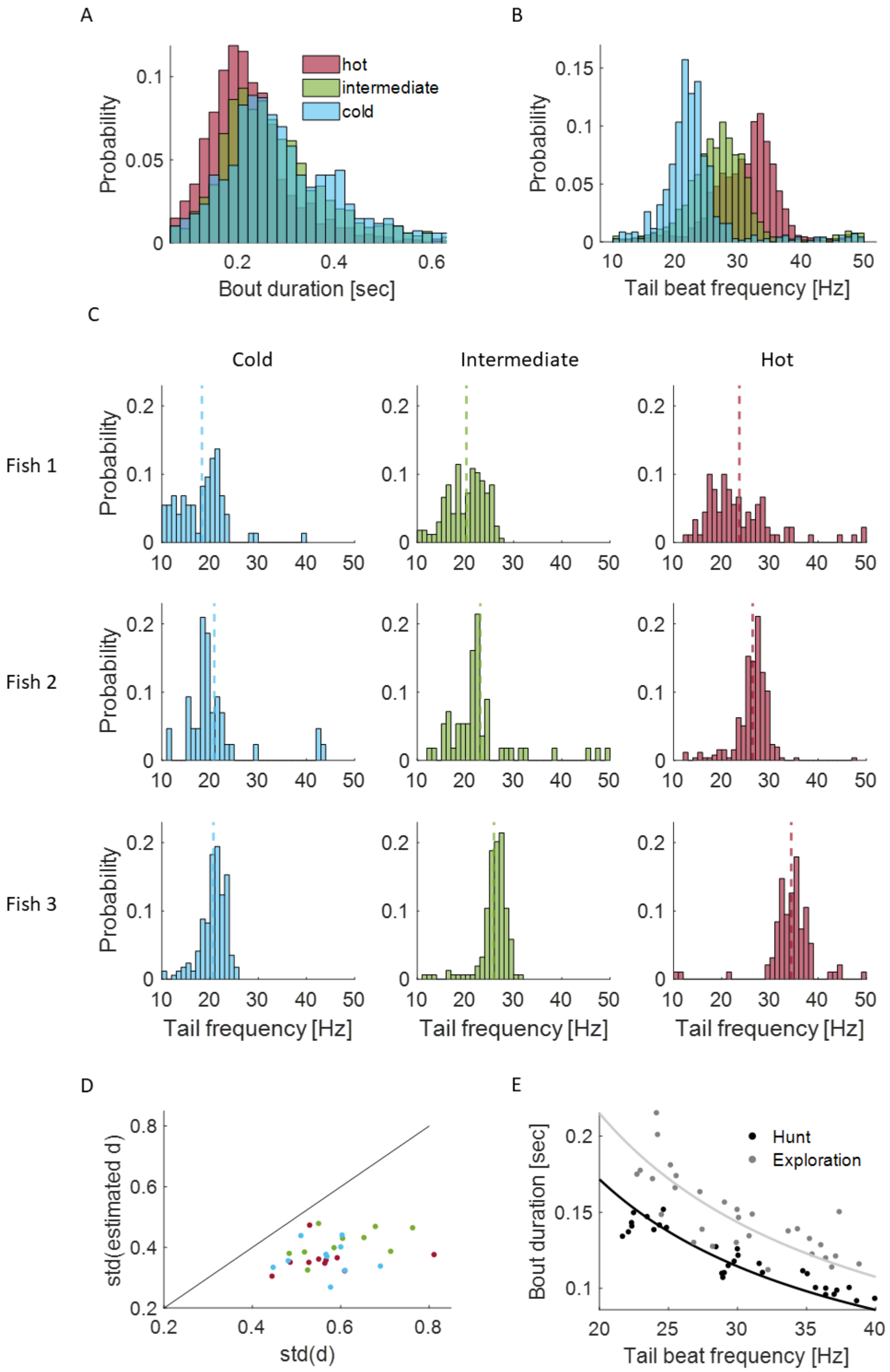


**Figure S6: Temperature modulates bout duration and tail beat frequency also in head-fixed and tail-free fish. (A)** Bout duration distributions differed significantly across temperature conditions in head-fixed, tail-free fish (Bonferroni-corrected KS-test, hot vs. intermediate $p = 10^{-34}$, intermediate vs. cold $p = 0.02$, hot vs. cold $p = 10^{-38}$). **(B)** Tail beat frequency significantly differed across temperatures in head-fixed, tail-free fish (Bonferroni-corrected KS-test, hot vs. intermediate $p = 10^{-98}$, intermediate vs. cold $p = 10^{-99}$, hot vs. cold $p = 10^{-235}$). **(C)** Tail beat frequency distributions of three example head-fixed fish across temperature conditions show inter-fish variability within each temperature, along with a consistent temperature-dependent shift. Dashed colored lines indicate the means. **(D)** The standard deviation of bout distance in freely swimming fish is lower than that of the estimated distance provided by the model across all fish and conditions. The black line is the unity line. **(E)** The relationship between tail beat frequency and bout duration. Assuming a constant distance ($d$) and varying frequency ($freq$), movement duration ($dur$) should be modulated accordingly: $freq \cdot dur \approx d$ and therefore $dur = d \cdot freq^{-1}$. Curves were fitted with a hyperbola for hunting and exploration (Matlab robust bi-square fit). The hyperbolic fit explains a greater proportion of variance for hunting ($R^2 = 0.74$) than exploration ($R^2 = 0.58$), indicating tighter coordination between frequency and duration in the hunting context.

**Movie S1: Temperature modulates larval zebrafish hunting behavior.**

Three hunting events recorded at three temperatures (left to right, cold, intermediate, hot), each with similar initial prey distance and angle relative to the fish. Recordings were captured from above at 500 fps and are displayed at 30 fps (slowed down by a factor of 16.67). Each video ends upon successful prey capture.

**Movie S2: Tectal neural responses to visual stimuli.**

Single-plane two-photon calcium imaging of a 14 dpf larval zebrafish at a depth of $70\mu m$, recorded at 29.946 fps and displayed at 180 fps (sped up by a factor of 6). The red dot indicates the visual stimulus. The recording was obtained under intermediate temperature condition ($27.5°C$).

**Movie S3: Behavioral recording of a head-fixed and tail-free fish.**

Tail movements of a head-fixed and tail-free fish were recorded at 500 fps and are displayed at 100 fps (slowed down by a factor of 5), showing a series of tail movements while neural activity is simultaneously recorded. The recording was obtained under intermediate temperature condition ($27.5°C$).
